# Amino acid repeat signatures underlying human-pathogen interactions

**DOI:** 10.1101/2025.03.17.643713

**Authors:** Anjali Kumari Singh, Nagashree Rachote, Anushka Agrawal, Vaidehi Sharma, Keertana Sai Kappagantula, Rajashekar Varma Kadumuri, Sreenivas Chavali

## Abstract

Emerging evidence suggests that amino acid homorepeats (HRs) in proteins (HRPs) contribute to protein interactability. What is the role of HRs in human-pathogen protein interactions? We find that pathogens engage physiologically important human HRPs, thereby affecting diverse host physiological processes. From the pathogen standpoint, (i) eukaryotic pathogens engage more HRPs but with host-sparse HRs, leading to disparate and discriminate interactions, (ii) prokaryotic pathogens engage less HRPs but with host-abundant non-polar HRs via host protein proxies bringing about discriminate or promiscuous interactions and (iii) viral pathogens engage more HRPs with host-abundant polar uncharged HRs affecting promiscuous interactions using host-partner HR tract mimicry. To propel further research, we introduce a resource Hi-PHI (http://hiphi.iisertirupati.ac.in/) cataloging critical information about human and pathogen HRPs and HRs. We propose mechanisms to (i) repurpose drugs targeting human HRPs engaged by pathogens for treating different infections and (ii) exploit HRs and their flanks as targets for pathogen-targeted anti-infectives.

## Introduction

Invasion of host by pathogens involves engaging host machinery effectively for the propagation and/or physiological functions of the pathogens, thereby triggering pathogenic response. This includes abrogating host physiological functions and/or hijacking host system. Pathogen proteins aid host invasion by (i) molecular mimicry to sequester host proteins and/or (ii) *de novo* pathological interactions to abrogate host physiological functions (eg. immune response), and/or hijack host machinery for pathophysiological outcomes (eg. viral replication)^1,2^. Pathogen proteins are known to interact with host proteins using protein functional units such as (i) protein domains, which are self-folding units that can function independently^3–6^ [**Figure S1A-S1B**] and (ii) sequence motifs, which are small stretches of consensus residues that are associated with molecular functions^7,8^ [**Figure S1C-S1D**].

Emerging evidence suggests that stretches of identical amino acids in proteins, referred as homorepeats (HRs), can act as interaction modules and contribute to high interactability leading to multifunctionality of proteins with HRs (HRPs)^9–11^. Importantly, human HRPs are associated with core processes, such as gene expression regulation, development and signalling in humans^9,10^, making them attractive targets for pathogens. On the other hand, the increased interaction potential conferred by HRs might be pivotal for pathogen proteins to better engage human proteins. How are human and pathogen HRPs and HRs engaged in human-pathogen interactions? In this study, we elucidate the role of HRPs and HRs in human-pathogen interactions by assembling, integrating and analysing diverse large-scale publicly available datasets pertaining to human-pathogen interactions, human functional and regulatory interactions, spatial gene-expression, phenotypic screens, drug-target networks and priority pathogens identified by World Health Organization (WHO) [**Figure 1A**].

**Figure 1.**
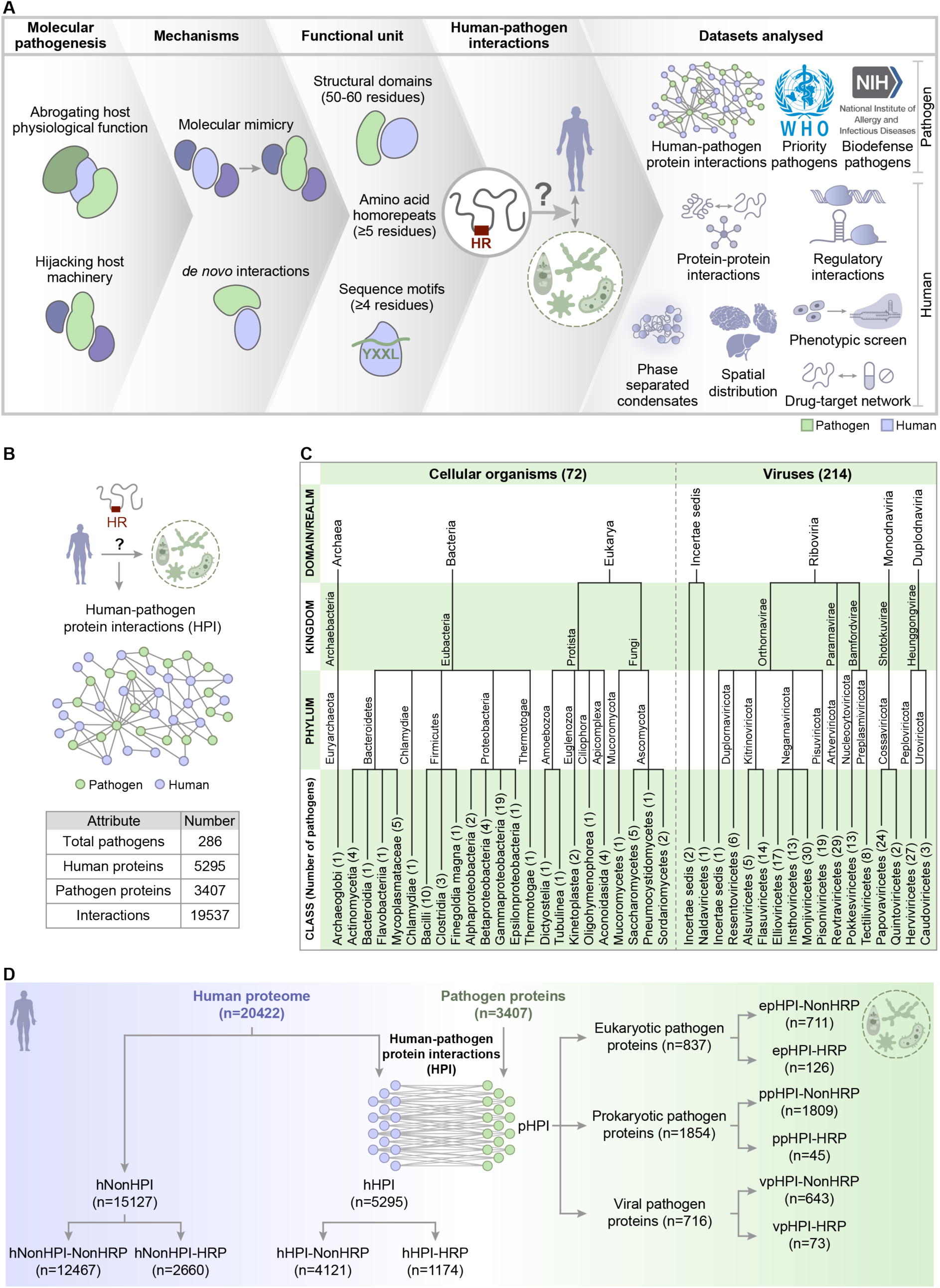
Modalities of human-pathogen protein interactions. **(A)** Illustration outlining the framework adopted in this study to understand the role of HRPs and HRs in human-pathogen pathophysiology. **(B)** Attributes of human-pathogen protein interaction (HPI) network assembled in this study. **(C)** Taxonomic classification of human unicellular and viral pathogens represented in HPI network. The numbers in parentheses indicate the number of species in each taxonomic class. **(D)** Classification of human and pathogen proteins, based on their participation in HPI and the presence of homorepeats (HR). n denotes the number of proteins in each class.

## Results

We assembled a comprehensive human-pathogen protein-protein interactome (HPI) comprising of 19,537 interactions between 5,295 human proteins and 3,407 pathogen proteins pertaining to a total of 286 human cellular (prokaryotic and eukaryotic) and viral pathogens spanning different taxa [**Figure 1B-1C**]. We classified proteins with identical amino acid runs (length ≥5) as HRPs, as previously defined^9,12^. Based on their participation in HPI and the presence of HRs, we classified the human proteins into HPI proteins with HRs (hHPI-HRP) and without HRs (hHPI-NonHRP), and those that do not participate in HPIs with HRs (hNonHPI-HRP) and without HRs (hNonHPI-NonHRP) [**Figure 1D**]. We drew comparisons for different attributes of hHPI-HRPs with hHPI-NonHRPs and hNonHPI-HRPs spanning phenotypes, molecular networks and gene-expression. Based on taxonomy and presence of HRs, pathogen proteins participating in the HPIs (pHPI) were categorized into those that belonged to (i) eukaryotic pathogens (epHPI-NonHRP and epHPI-HRP), (ii) prokaryotic pathogens (ppHPI-NonHRP and ppHPI-HRP) and (iii) viral pathogens (vpHPI-NonHRP and vpHPI-HRP). To understand the taxa-specific attributes of pathogen HRPs, comparisons were drawn between (i) the HRPs and NonHRPs belonging to each of the three taxa and (ii) the pathogen HRs across the three taxa [**Figure 1D**].

## Physiologically important human proteins with homorepeats are engaged by pathogens

Are human HRPs engaged in HPI? Human HRPs, especially those engaged by prokaryotic and viral pathogens, are enriched among human proteins that participate in HPI [**Figure 2A-2B**]. Importantly, we find that hHPI-HRPs engaged by prokaryotic and viral pathogens are predominantly enriched for regulatory processes such as chromatin organization and transcription, post-transcriptional regulation [**Figure 2C**]. Additionally, hHPI-HRP partners of prokaryotes and viruses are also enriched for development and differentiation, while those of the prokaryotes are enriched for signalling. We could not find any enriched biological processes for the hHPI-HRPs engaged by eukaryotic pathogens. This can be due to (i) fewer hHRPs identified thus far to be interacting with eukaryotic pathogens and/or (ii) the hHRPs interacting with eukaryotic pathogen proteins could be participating in diverse biological processes. Contrarily, hHPI-NonHRPs are predominantly enriched for post-translational regulation, cell death and signalling across the three taxa of pathogens [**Figure 2C**]. Jaccard similarity indices across different biological processes show that hHPI-NonHRP interacting partners of prokaryotic and viral pathogen proteins are more shared across biological processes than the hHPI-HRPs [**Figure 2D**]. This suggests that prokaryotic and viral pathogens target hHPI-HRPs that participate/regulate specific processes.

**Figure 2.**
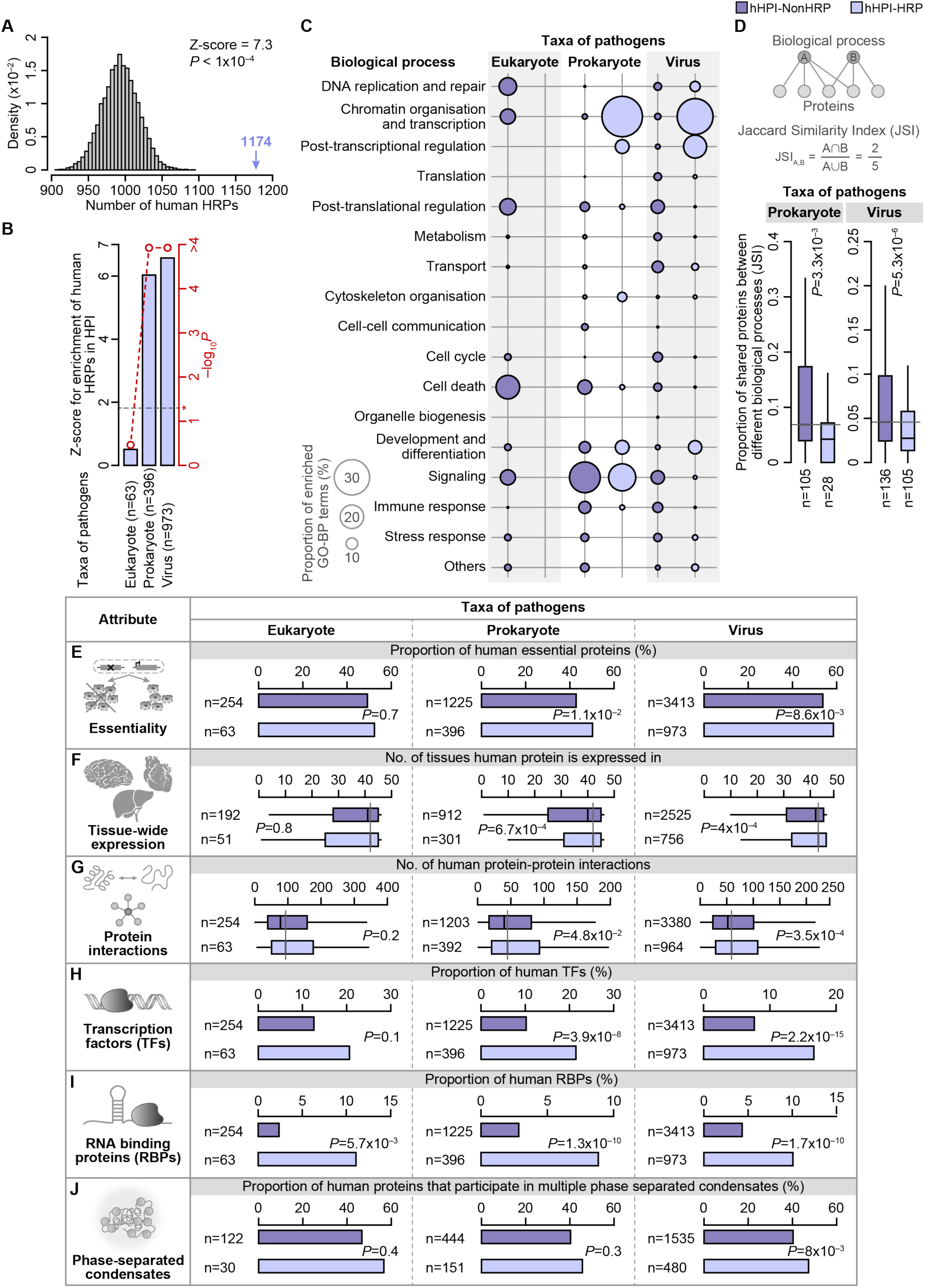
Physiological importance of human proteins engaged by pathogens. (**A**) Enrichment of HRPs in human proteins that are engaged by pathogens. The histogram in grey represents the random expectation and the blue arrow denotes the actual observation of the number of hHPI-HRPs. **(B)** Enrichment of human HRPs that are engaged by different taxa of pathogens. The dotted red line denotes the significance estimate and the dotted black line highlighted by the asterisk represents the threshold for statistical significance. Z-scores and P values were estimated using permutation testing. **(C)** Bubble plot indicating the proportion of significantly enriched (FDR<0.05) Gene Ontology Biological Process (GO-BP) terms in each of the manually classified major biological processes. The size of the bubble indicates the proportion of enriched GO-BP terms in each broad category for a given protein class in each taxon. **(D)** Proportion of shared proteins across different biological processes, estimated using Jaccard Similarity Index (JSI). n denotes the number of pairs in each class. P value was estimated using Wilcoxon rank sum test. Bar plots showing the proportion of human **(E)** essential proteins, **(H)** transcription factors (TFs), **(I)** RNA binding proteins (RBPs) and **(J)** proteins that participate in more than one human phase separated condensates, that are engaged by different pathogens in HPI. P values were computed using Fisher’s Exact test. Box plots representing the **(F)** total number of tissues in which the different classes of human proteins are expressed and **(G)** number of protein-protein interactions in humans. P value was computed using Wilcoxon rank sum test. n denotes the total number of proteins in each class.

What is the physiological relevance of hHPI-HRPs? Physiological importance can be assessed through various attributes such as gene essentiality, tissue-wide expression, centrality in human protein-protein interaction network and/or impact on global regulation. We find that hHPI-HRPs engaged by prokaryotic and viral pathogens are (i) preponderantly essential, (ii) have broader tissue distribution in humans, (iii) have higher number of protein-protein interactions and (iv) enriched for human regulatory HRPs involved in transcriptional and post-transcriptional regulation [**Figure 2E-2I**]. Strikingly, hHPI-HRPs engaged by viral pathogens predominantly participate in multiple biological condensates, which represent both higher order assemblies resulting from multiple interactions as well as dynamic regulatory units [**Figure 2J**]. The attributes that represent physiological importance are lot more pronounced in hHPI-HRPs compared to hHRPs that do not participate in HPI (hNonHPI-HRP) [**Figure S2**]. Collectively, these findings imply that functionally important human HRPs are engaged by pathogens in HPI, affecting a diverse range of host biological processes, including affecting global regulation across tissues.

## Sequestration of hHRPs by pathogens might significantly impact human physiology

How important are hHPI-HRPs for human physiology? To mimic the impact of sequestration of human proteins by pathogen proteins on human protein interaction network, we performed network dismantling. For this, we removed each protein (node) in the human protein-protein interaction network, corresponding to a virtual knock-out of that protein, and then assessed for the impact of node removal on the network topology by estimating the (i) link density– average global connectivity of the network^13^ and (ii) assortativity– preferential association of proteins with similar interaction potential^14^ [**Figure 3A-3C**]. Removal of hHPI-HRPs that interact with prokaryotic and viral pathogens significantly (i) reduces the link density, resulting in sparsely connected networks and (ii) enhances the assortativity, leading to an increase in disjoint modules, compared to removal of hHPI-NonHRPs [**Figure 3D**]. While hHPI-HRPs participate in limited number of processes [**Figure 2C**], they exhibit high interactability and inter-connectivity in human protein interaction network [**Figure 3C**] implying that they might connect pathways or protein complexes involved in closely related biological processes. These observations suggest that the hHPI-HRPs engaged by prokaryotic and viral pathogen proteins bring about more interactability at an individual protein-level, denser connectivity at the network-level, facilitating inter-connectedness across modules affecting fundamental processes.

**Figure 3.**
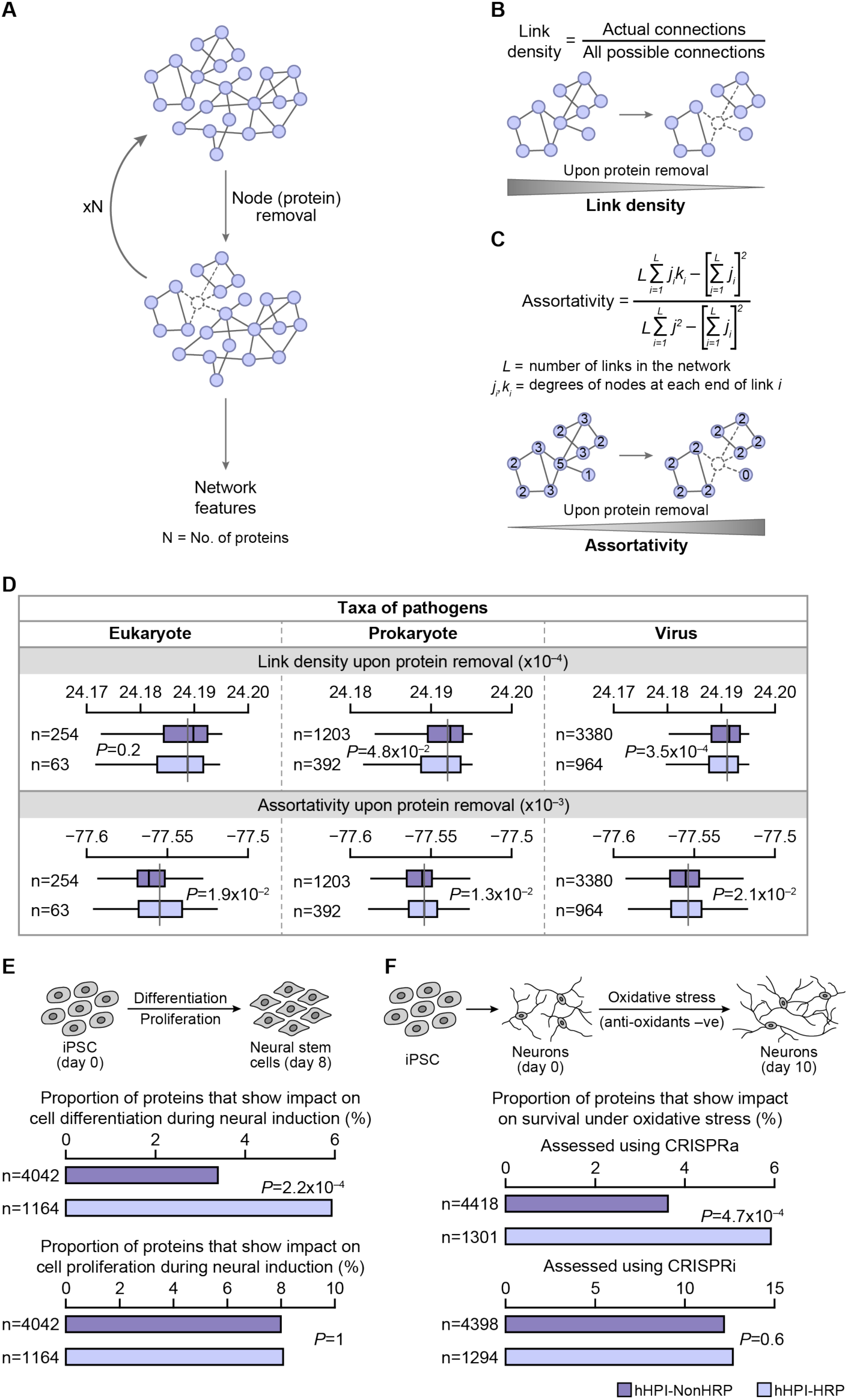
Impact of human HRPs engaged by pathogens on human protein-protein interaction network and on different human cellular phenotypes. Schema illustrating **(A)** the network dismantling approach, and estimation of the network topological attributes– **(B)** link density^13^ and **(C)** assortativity^14^ upon node (protein) removal from the human protein-protein interaction network. The numbers in the nodes represent the number of connections (degree) for each node. **(D)** Box plots representing the link density (top panel) and assortativity (bottom panel) of human proteins that are engaged in HPIs. P value was computed using Wilcoxon rank sum test. Bar plots showing the proportion of proteins that show impact on **(E)** cell differentiation and cell proliferation during neural induction of human iPSCs and **(F)** survival of neurons in oxidative stress assessed using CRISPRa and CRISPRi. n denotes the total number of proteins in each class. P value was estimated using Fisher’s exact test.

To decipher the plausible phenotypic impact when hHPI-HRPs are engaged by pathogens, we investigated publicly available genome-wide CRISPRi/a-based phenotypic screens assessing for (i) cell fate determination, (ii) cell proliferation and (iii) cell survival during stress. Compared to HPI-NonHRPs and NonHPI-HRPs, removal of hHPI-HRPs tend to significantly influence the neural differentiation of human induced pluripotent stem cells (iPSC) but not cell proliferation [**Figure 3E**; **Figure S2H**]. This suggests that the hHPI-HRPs are key for neuronal differentiation. While, the impact of removal of hHPI-HRPs is comparable to that of hHPI-NonHRPs for neuronal survival under oxidative stress [**Figure S2I**], there is a significant change on the survival of neurons when hHPI-HRPs are activated using CRISPRa [**Figure 3F**]. This finding suggests that upregulation of hHPI-HRPs influences stress response behaviour of neurons in a dosage-dependent manner. Thus, engaging such proteins by pathogens would have greater impact on host physiology.

## Pathogens of different taxa show diverse modes of engaging pHRPs

How do pathogens employ their HRPs to engage human proteins? Analysis of the number of pathogen HRPs (pHRPs) of different taxa revealed that higher proportion of HRPs of eukaryotic pathogens (epHPI-HRPs) participate in HPI, compared to those of prokaryotes and viruses [**Figure 4A**]. Although more in number, epHPI-HRPs predominantly engage in solitary interactions with human proteins, bringing about one-to-one or discriminate interactions [**Figure 4B-4C**]. Contrarily, HRPs of prokaryotic pathogens (ppHPI-HRPs) bring about discriminate as well as one-to-few/many i.e. promiscuous interactions with human proteins [**Figure 4B-4C**], while HRPs of viral pathogens (vpHPI-HRPs) predominantly engage in promiscuous interactions with human proteins [**Figure 4B-4C**]. These findings capture considerable diversity in pHRP deployment in terms of number of pHRPs and their mode of engagement of human proteins across different taxa of pathogens.

**Figure 4.**
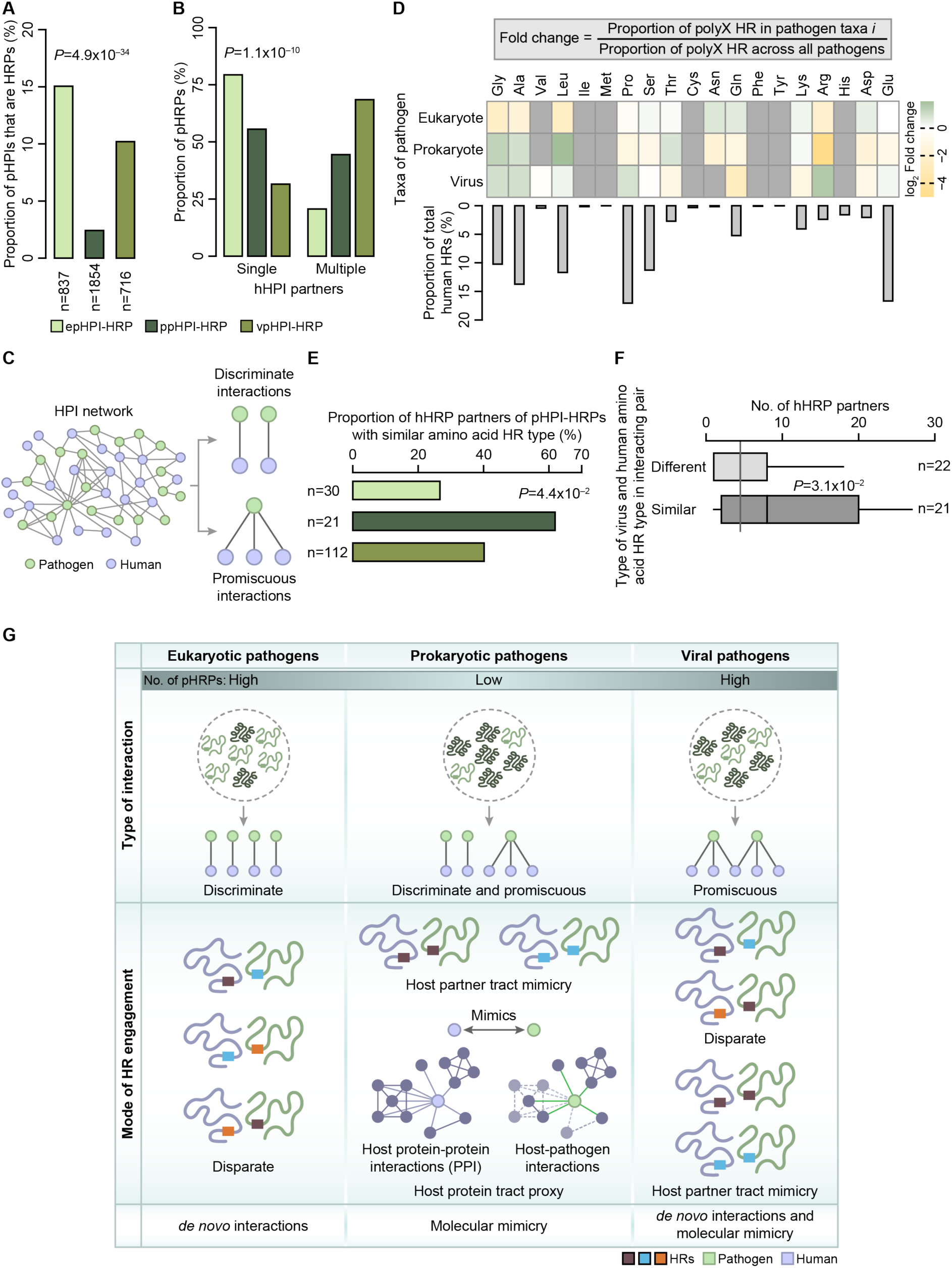
Modes of HRP and HR engagement by different taxa of human pathogens. Bar plot showing the proportion of pHRPs **(A)** belonging to pathogens of different taxa (ep-eukaryotic pathogens, pp-prokaryotic pathogens and vp-viral pathogens) that participate in HPI, with n denoting the number of proteins in each class, and **(B)** that engage single or multiple human proteins in HPIs. P value was estimated using Fisher’s Exact test. **(C)** Illustration depicting different modes by which pHPI-HRPs engage human proteins in HPI. **(D)** Heatmap showing fold change of different amino acid HR types of different taxa of pathogens that are engaged in HPI over the total HRs engaged in HPI (top panel). Bar plot showing the distribution of HR type in human proteome (bottom panel). **(E)** Bar plot showing the proportion of interactions of pHRPs and hHRPs with similar amino acid HR type. n denotes the number of unique interactions in each taxon. P value was estimated using Fisher’s Exact test. **(F)** Boxplot depicting the total number of human interactors of viral HRPs with at least one human HRP interactor with similar amino acid HR type (similar) and viral HRPs with human HRP interactors with different amino acid HR types (different). n denotes the number of unique viral HRPs in each class. P value was estimated using Wilcoxon rank sum test. **(G)** Illustration summarizing the different modes of HRP and amino acid HR type engagement by human pathogens of different taxa.

## Amino acid HR type might influence diverse modes of interactions of pHRPs across different taxa

What is the impact of the nature of amino acid HRs on the variability in the mode of interactions of pHRPs belonging to different taxa of pathogens? To address this, we examined the proportion of different amino acid pHRPs across the three different taxa in HPIs. We found that HRPs of eukaryotic pathogens (epHPI-HRPs) show a higher tendency for the presence of polar uncharged HRs such as polySer, polyThr, polyAsn and polyGln [**Figure 4D**]. Strikingly, human proteome is depleted for these HR types (except polySer) [**Figure 4D**]. Furthermore, epHPI-HRPs interact with human HRPs that lack HRs with similar physicochemical properties [**Figure 4E**]. For example, the epHRP serine/threonine-protein phosphatase PPZ1 (polyGlu and polyGln containing HRP) of *Candida albicans* interacts with human cell cycle protein serine/threonine-protein phosphatase 1 regulatory subunit 10 PPP1R10 (polyGly, polyLys and polyPro containing hHRP) and cytoskeletal protein neurabin-2 PPP1R9B (polyPro containing hHRP)^15^. This implies that eukaryotic pathogens employ HRs that are less-frequent in the human proteome and engage the human proteins using “host-sparse” HRs probably leading to *de novo* interactions. This type of HR engagement could lead to disparate discriminate interactions [**Figure 4G**].

Conversely, HRPs of prokaryotic pathogens (ppHRPs) show an overrepresentation of non-polar HRs such as polyGly, polyAla and polyLeu [**Figure 4D**]. Strikingly, (i) such amino acid HR types are predominant in the human proteome and (ii) the hHPI-HRPs that interact with ppHPI-HRPs have amino acid HR types of similar physicochemical nature [**Figure 4E**]. For instance (i) the cell division protein FtsZ (polyGly containing ppHRP) protein in *Francisella tularensis* subsp. *tularensis* SCHU S4 interacts with human proteins involved in cell adhesion and cytoskeleton proteins including protocadherin-16 DCHS1 (polyLeu containing hHRP), phospholipid scramblase 1 (PLSCR1) and moesin (MSN) as well as transcription factors involved in immune response TNFAIP3-interacting protein 1 TNIP1^16^ and (ii) inner membrane permease of ABC transporter fhuB of *Yesinia pestis* (polyLeu containing ppHRP) interacts with human transcription factors, immune response and cell proliferation proteins including nuclear factor NF-κB p105 subunit (NFKB1; polyGly containing hHRP), cyclic AMP-dependent transcription factor ATF-6α (ATF6) and thioredoxin reductase 1 (TXNRD1)^16^. This implies that HRPs of prokaryotic pathogens engage human HRPs by (i) mimicking the host partner tract or (ii) hijacking host protein neighborhood through host protein proxy, specifically non-polar HR containing HRPs via hydrophobic interactions facilitating promiscuous interactions [**Figure 4G; Figure S3**].

Alternatively, viral pathogen HRPs (vpHRPs) engage HRs such as polyGly, polyAla, polyPro and polyArg to engage host proteins [**Figure 4D**], which are also predominant in the human proteome (except polyArg). The interactions between vpHRPs and human HRPs containing both similar (mainly non-polar HR) as well as different HR types are comparable [**Figure 4E**]. Interestingly, compared to those that interact with different HR type containing human HRPs vpHRPs that interact with similar HR tract containing human HRPs have higher number of host interactions probably contributing to promiscuous interactions [**Figure 4F**]. For instance, polyPro-containing UL27 vpHRP of Human herpesvirus 5 (strain Merlin) interacts with multiple non-polar HR-containing human proteins– 26S proteasome non-ATPase regulatory subunit 3 (PSMD3; polyAla and polyPro HRs), E3 ubiquitin-protein ligase UBR5 (UBR5; polyAla, polyLeu and polySer HRs), WD repeat-containing protein 26 (WDR26; polyGly and polySer HRs), protein disulfide-isomerase A4 (PDIA4; polyGlu and polyLeu HRs), E1A-binding protein p400 (EP400; polyGlu, polyPro and polyGln HRs) and transformation/transcription domain-associated protein (TRRAP; polyPro HR)^17^. This implies that viral pathogen HRPs show both disparate interactions leading to *de novo* interactions as well as interactions using host-partner tract mimicry, the latter bringing about promiscuous interactions [**Figure 4G**]. Collectively, these findings highlight that pathogens belonging to different taxa show distinct patterns in employing different amino acid HR types to engage human proteins.

## Discussion

Systems-level and molecular-level studies so far have elucidated the role of protein domains and sequence motifs in host-pathogen interactions^6,8,18^. However, little is known about how amino acid repeats, an emerging class of interaction modules, are engaged in host-pathogen interactions. Here, we present a comprehensive systems-level analyses and show how different taxa of pathogens engage both human and pathogen proteins with homorepeats distinctly in human-pathogen protein interactions. We find that (i) HRPs are engaged by both human and pathogens during host-pathogen interactions and (ii) pathogens engage physiologically relevant human HRPs in HPI, whose depletion might have severe consequences to host physiology and (iii) pathogens across different taxa engage pHRPs of diverse HR types differently in HPIs, bringing about distinct modes of engagement of host proteins.

Notably, some of the priority pathogens identified by World Health Organization (WHO) based on parameters such as mortality, incidence, non-fatal health burden, transmissibility, preventability and treatability such as *Mycobacterium tuberculosis*, *Escherichia coli*, *Candida parapsilosis* and *Candida tropicalis* engage high number of pHRPs in HPIs and/or have high HRP content in their proteomes [**Figure S4**]. Similarly, some of the biodefense pathogens classified by National Institute of Allergy and Infectious Diseases (NIAID) owing to their ability to cause diseases of high public health concern such as *M. tuberculosis*, *Escherichia coli*, *Bacillus anthracis* and *Yersinia pestis* engage relatively higher number of pHRPs in HPIs [**Figure S4**]. Strikingly, pathogens causing life-threatening diseases such as malaria (*Plasmodium falciparum*; causing nearly 250 million cases and 600,000 deaths worldwide in 2022), and tuberculosis (*M. tuberculosis* causing ∼1.3 million deaths in 2022) which also results from multi-drug resistance, engage more pHRPs in HPIs. These findings imply that high-risk human pathogens show higher number of HRPs in their proteomes and/or employ more number of pHRPs in HPIs. From a viral pathogen perspective, (i) viruses such as human herpesvirus 5 and Epstein-barr virus that infect multiple organs and organ systems, and/or (ii) epidemic– and pandemic-causing life-threatening viruses such as Ebola and SARS coronavirus have high HRPs in their proteomes and/or engage more HRPs in HPIs [**Figure S4**]. Furthermore, in addition to the presence of HRs, repeat-associated variation can also contribute to pathogenicity. This is best exemplified by the interaction of polyHis HR of Knob-associated histidine-rich protein (KAHRP) in *P. falciparum* with the β-spectrin of human infected erythrocytes in a repeat length-dependent manner, affecting cytoadhesion and hence the pathogenesis^19–21^. These facets emphasize the need for immediate and thorough investigations to delineate the impact of different amino acid HRs and repeat-associated variation on pathogenicity and virulence.

From the pharmacological stand point, reconstruction of human-drug interaction network of hHPI-HRPs revealed that many HRPs are already being targeted to alleviate certain infectious diseases [**Figure 5A**]. For instance, (i) human nuclear factor NF-kappa-B NFκB1 with polyGly HR is targeted in HIV, herpes simplex virus as well as SARS CoV-2 infections^22,23^. In the HPI network, human NFκB1 interacts with 8 different bacterial and viral pathogens, amounting to about 172 human-pathogen protein interactions, (ii) human epidermal growth factor receptor (EGFR) is targeted in several skin and soft tissue infections^24^. In the HPI network, human EGFR, which contains polyLeu HR, interacts with 8 different bacterial and viral pathogens, with 19 pathogen protein interactions, and (iii) human plasminogen (PLG), which contains polyLeu HR, is targeted by drugs for conjunctivitis^25^ interacts with 6 eukaryotic and bacterial pathogens, participating in a total of 21 human-pathogen protein interactions. These instances highlight that some of the existing drugs that target these important hHRPs to treat few infectious conditions, can be repurposed for treating and/or alleviating symptoms of other infections, caused by pathogens even from different taxa.

**Figure 5.**
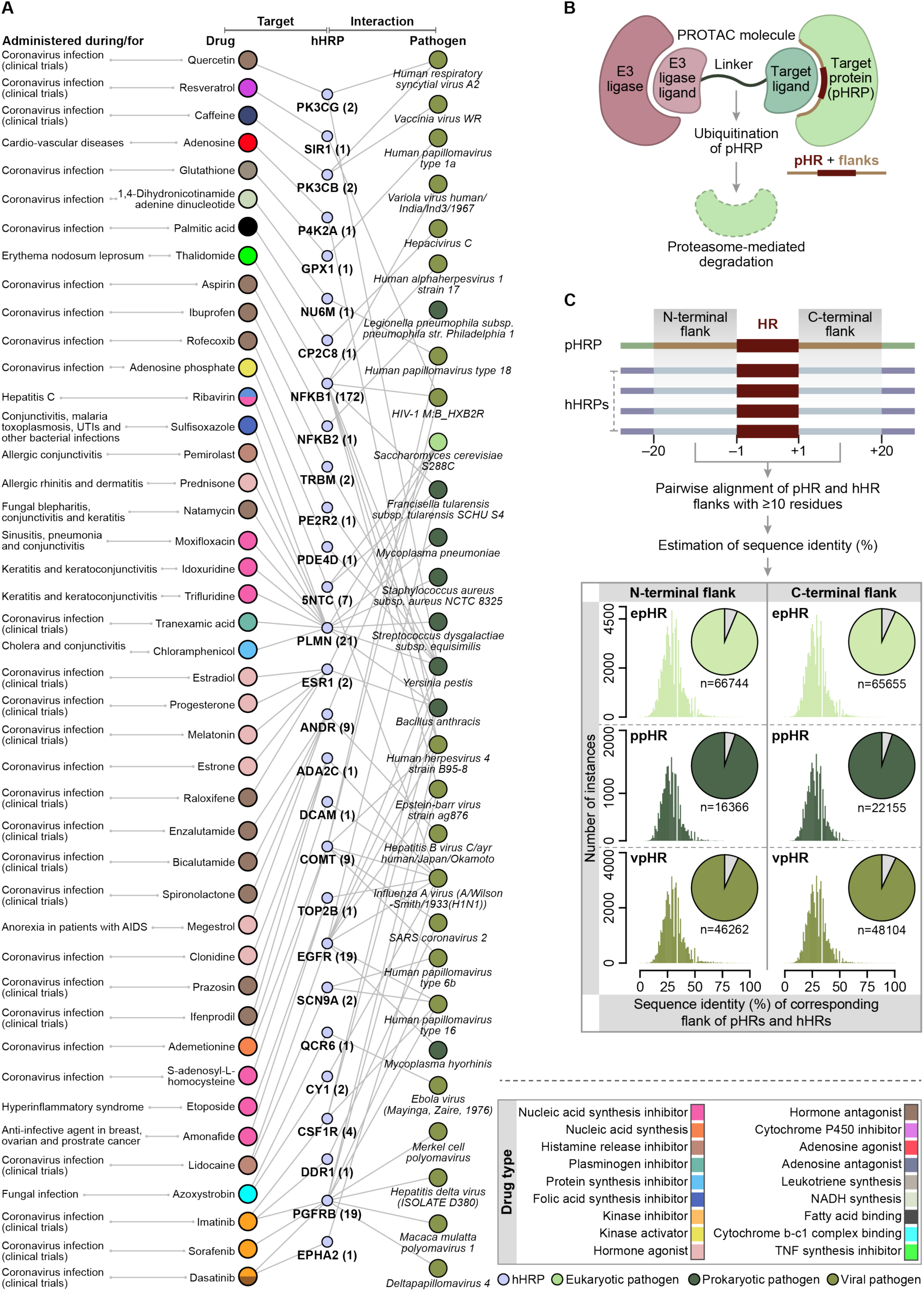
Therapeutic intervention strategies for targeting human and/or pathogen HRPs or HRs that participate in HPI. **(A)** Tripartite network with connections between FDA-approved drugs/molecules under clinical trial administered during infections (left nodes), drug target hHRPs participating in HPI (middle nodes) and pathogens that engage the hHRPs identified in this study (right nodes). The colour of the drug nodes denote the drug type based on function, while that of the pathogen nodes on the right denote the taxonomic class of the pathogen. The number in parenthesis for hHRP nodes indicates the total number of pathogen protein partners of each hHRP. The conditions for which the drugs are being currently administered are provided on the left side of the drugs. **(B)** Schema representing the approach that involves HR– and their flanks-aided PROTACs-based protein targeting and degradation. **(C)** Pairwise alignment of the N-terminal and C-terminal flanking residues of pathogen HRs with corresponding flanks of the same amino acid HR type in human proteome (top panel). Histograms showing the sequence identity of HR-associated flanks of pathogen and human proteins with same amino acid HR type (bottom panel). The pie chart shows the total pairs of pathogen and human flanking sequences that qualify (lighter colour in the pie) for the analysis (at least 10 residues from pathogen and human flanking sequences should be aligned). The grey part of the pie denotes pathogen HRs and the flanks, with <10 residues aligning with the regions in human proteome, are ideal for designing species-specific PROTAC molecule without any cross-reactivity with human proteins. n denotes the total number of pathogen and human HR-flanking pairs that were analysed for the study.

Additionally, some of the human HRPs can be also be targeted for host-directed therapies using alternate approaches. For instance, human HRs along with their flanks can be designated targets to abrogate human-pathogen interactions. Especially in case of host-partner tract mimicry of human HRs by pathogens, HR-based targeting can be used as a potential therapeutic intervention strategy. Similarly, pathogen-directed therapies especially for eukaryotic pathogens that bring about disparate interactions with human proteins by engaging host-sparse HRs can also be developed, with minimal impact on host system. One approach to achieve this, is to employ a targeted protein degradation method using proteolysis-targeting chimeras (PROTACs)^26,27^ which utilizes the ubiquitin-proteasome system to degrade a target protein [**Figure 5B**]. We observed that the sequences flanking the repeats on either side of pathogen proteins show very less similarity with human proteins [**Figure 5C**]. This makes such unique sequences which include the HRs and the flanks ideal candidates to be targeted by PROTACs with high specificity and low cross-reactivity with host proteins. To propel research to investigate the role of HRs in pathogenicity and virulence of human infectious agents we have developed a database named Hi-PHI (<u>H</u>omorepeats in <u>P</u>athogen-<u>H</u>uman Interactions; http://hiphi.iisertirupati.ac.in/) which catalogs all the data generated and analysed in this study [**Figure S5**]. Furthermore, Hi-PHI also provides percentage sequence identity of pathogen HRs along with their flanks with the HRs and flanks in the entire human proteome [**Figure 5C**], to facilitate designing pathogen-directed targeting approaches. Notably, large pharmacological screens such as those identifying epitopes for widely used antibodies for disease treatment or to predict cross-reactivity^28–31^ often do not include HRs, may be due to the inherent difficulties associated with studying such low-complexity sequences. We anticipate that the results and the datasets provided here will aid in designing studies that target human HRs and/or pathogen HRs, to abrogate host-pathogen interactions, thereby attenuating infections.

## Limitations of the study

The findings in this study are contingent to the large-scale datasets integrated and analyzed, which might influence some of the interpretations drawn here. This includes the number of human-pathogen protein interactions and the density of interactions per pathogen identified thus far. Furthermore, through systems-level analyses, we have obtained general trends and every trend reported might not be applicable to each protein in every species across the different taxa studied here. While the network dismantling and phenotypic screen analyses provide important insights, they essentially do not present an exhaustive picture of the impact of engagement of hHRPs by pathogens on human physiology. Although, this study provides a comprehensive account of the different modes of HRP-engagement in HPI and potential roles of HRs, targeted experimental investigations are required to elucidate the functional roles of both hHRs and pHRs.

## Acknowledgements

We thank P.L. Chavali, A. Dhayalan, and A.D. Allu for their feedback on the manuscript. This study was supported by IISER Tirupati core funding (A.A., V.S., R.V.K., and S.C.); Prime Minister’s Research Fellowship, Government of India (A.K.S., N.R.); Ramalingaswami Re-entry Fellowship BT/RLF/Re-entry/05/2018 from Department of Biotechnology, Government of India (R.V.K., and S.C.); Junior/Senior Research Fellowships from Department of Biotechnology, Government of India (K.S.K.); Core Research Grant CRG/2023/004691 from Science and Engineering Research Board, Government of India (A.K.S, and S.C.).

## Author contributions

Conceptualization: SC; Methodology: SC, AKS; Formal analysis: AKS, NR, AA, VS, KSK; Visualization: AKS, NR, RVK, SC; Writing—original draft: SC, AKS; Writing—review & editing: SC, AKS, NR, AA, VS, KSK, RVK; Supervision: SC

## Materials and Methods

### Assembly of human-pathogen interactome

Human-pathogen protein interactions (HPI) were assembled from HPIDB 3.0^32,33^, MorCVD^34^, HoPaCI-DB^35^, BIOGRID^36^ and high-throughput study^37^. The interactions were mapped to subset one strain per pathogen species (except for viruses), and then mapped to unique UniProt identifiers. HPIs obtained using yeast-2-hybrid system were disregarded. The human and pathogen protein sequences were retrieved from UniProt. Pathogen taxonomy information was extracted from NCBI Taxonomy browser^38^. An in-house Perl script was used to identify homorepeats (HR; amino acid type, length and location in the protein) in the human and pathogen proteins. A protein was considered as an HRP if it had at least one HR stretch with a minimum length of 5 residues, as previously defined^9,12^.

### Gene ontology enrichment analysis

Assessment of enriched Gene Ontology-Biological Processes (GO-BP) was done by DAVID server using an FDR cut-off of < 0.05^39,40^. The enriched GO-BP terms were then manually organized into major biolgical processes. We computed the extent of overlapping genes between any two enriched processes using Jaccard Similarity Index, which represents the ratio of the number of genes common to both biological processes to the sum of all genes in the two processes, as previously described^12^.

### Assembly of human proteome-level datasets

Dataset for human (i) essential genes/proteins, (ii) tissue-wide expression of proteins, (iii) physical protein-protein interaction network, (iv) protein complexes, (v) transcription factors and RNA binding proteins and (vi) phase-separated condensates were obtained from previously published study^12^. Human drug-target network for hHPI proteins was reconstructed using data obtained from the drug-target network^25^ and STITCH database^41^. Drug-target interactions from STITCH with binding data were considered here. The dataset was curated for drugs that target hHPI proteins involved in diseases caused by cellular and viral pathogens. The curated dataset comprised of 251 human proteins which were targets of 179 drugs. The functional classification of drugs based on their mode of action was obtained from DrugBank^42^.

### Assembly of pathogen-related datasets

We compiled the disease caused by each pathogen and the number of affected organs and organ systems through extensive literature search. The information for viral pathogens was curated from Human Virus Database (HVD)^43^. We obtained a list of 20 bacterial (in 2024) and 19 fungal (in 2022) priority pathogens identified by the World Health Organization (WHO; https://www.who.int/). This categorization of priority pathogens is based on parameters such as mortality, incidence, non-fatal health burden, transmissibility, preventability and treatability. We also obtained the list of biodefense pathogens, those that cause diseases of high public health concern, from National Institute of Allergy and Infectious Diseases (NIAID; https://www.niaid.nih.gov/).

### Network analysis

We computed the topological properties of human-pathogen protein interaction network and human protein-protein interaction network using Cytoscape 3.0^44^ and in-house Python and R scripts. Using an in-house written python script, we performed network dismantling of human protein-protein interaction network, by removing one node (protein) at a time. This led to the generation of 17,144 networks, each of which lacked one human protein. To assess the impact of a protein on protein interaction network, we estimated the network topological properties such as link density and assortativity for the network resulting from the removal of that protein and compared the network properties of the real network.

### Assembly of large-scale phenotypic screens

We assembled (i) CRISPRi-mediated genome-wide phenotypic screen comprising of 18,489 human coding genes involved in cell proliferation and differentiation during neural induction of human iPSCs to form neural stem cells from Wu et al^45^, and (ii) CRISPRa-mediated phenotypic screen comprising of 20,115 human genes and CRISPRi-mediated phenotypic screen comprising of 20,071 human genes involved in survival of neurons under oxidative stress (without anti-oxidant treatment) from Tian et al^46^.

### Estimation of percentage sequence identity

We retrieved 20 residue flanking the N-terminus and C-terminus of each HR in the human and pathogen proteins. This resulted in 5,412 N-terminal and 5,963 C-terminal flank sequences of hHRs and 342 N-terminal and 332 C-terminal flank sequences of pHRs. We disregarded flanks with <20 residues. For each flanking sequence of a pathogen HR, we computed pairwise percentage sequence identity using an in-house written R script (defined as the ratio of number of aligned residues to total length of the alignment) with corresponding flank of all human proteins with the same amino acid HR type. For instance, pair-wise percentage sequence identity of the N-terminal flank of a pathogen polyGly HR was estimated with the N-terminal flank of all human polyGly HRs. We could not perform pair-wise alignments of flanks with <10 aligning residues from either pathogen or human proteins. This led to 117,846 (out of 129,372) and 122,978 (out of 135,914) pairwise alignments of N-terminal and C-terminal flanks, respectively.

### Statistical analysis

Statistical significance for the differences in the distribution of discrete variables was assessed using Fisher’s Exact test and that of continuous variables was estimated using the non-parametric Wilcoxon rank sum test. Enrichment of proteins in different classes (for instance, human HRPs in HPI) was examined using permutation tests by performing 10,000 randomizations. In each permutation, the protein of interest (e.g., human proteins that participate in HPI) was replaced with a random protein and the number of random proteins that overlapped with a specific class of proteins (e.g., human HRPs) was noted for all the randomizations. We estimated Z score and P value from the distribution of overlapping proteins from the random expectations. Z score represents the magnitude of deviation of the actual observation compared to the mean of random distribution in terms of the number of standard deviations. P values for enrichment were estimated as the ratios of the number of the randomly observed proteins ≥ to the number of actually observed proteins to the total number of randomized samples (10,000). All statistical analyses were performed in R.

