## Supplemental Information for "Amino acid repeat signatures underlying human-pathogen interactions"

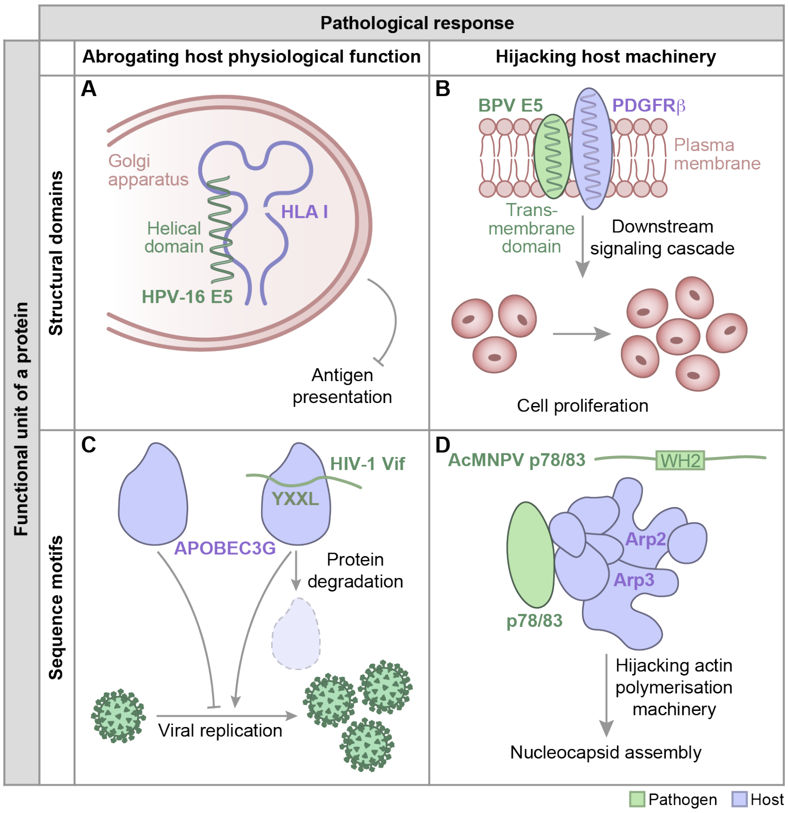


**Figure S1. Engagement of human proteins by pathogens using protein domains and sequence motifs. (A)** Human papillomavirus type 16 E5 protein (HPV-16 E5) prevents transport of the major histocompatibility class I (MHC I; HLA class I in humans) to the cell surface by interacting with HLA-I heavy chain by the first hydrophobic helical domain of E5, interfering with antigen presentation and abrogating host immune response[1]. **(B)** Small transmembrane protein E5 of bovine papillomavirus (BPV) forms a stable complex in dimerized form with PDGFRb proteins, via transmembrane domains, causing receptor dimerization, which activates the receptor and subsequently triggers downstream mitogenic signalling, enhanced DNA synthesis and cell transformation, leading to cancers[2-4]. **(C)** Human immunodeficiency virus-1 (HIV-1) accessory protein Vif binds to human APOBEC3G, involved in restriction of viral replication, via $_{69}$YXXL$_{72}$ motif, in HIV-1-infected CD4 T$^{+}$ cells. This binding targets human A3G for polyubiquitination and degradation, thereby leading to attenuation of host-defense response[5]. **(D)** Viral capsid protein p78/83 of Autographa californica M nucleopolyhedrovirus (AcMNPV) contains the WH2 (WASP homology) motifs (AcMNPV1: $_{225}$DDRQQLLEAIRNEKN–RTR$_{242}$; AcMNPV2: $_{321}$PSLSNVLSELKSGTV–RLK$_{338}$). This motif possibly facilitates binding to and activation of the Arp2/3 complex, which allows the virus to engage the host-cell actin cytoskeleton machinery for nucleocytoplasmic transport and assembly of nucleocapsid[6].

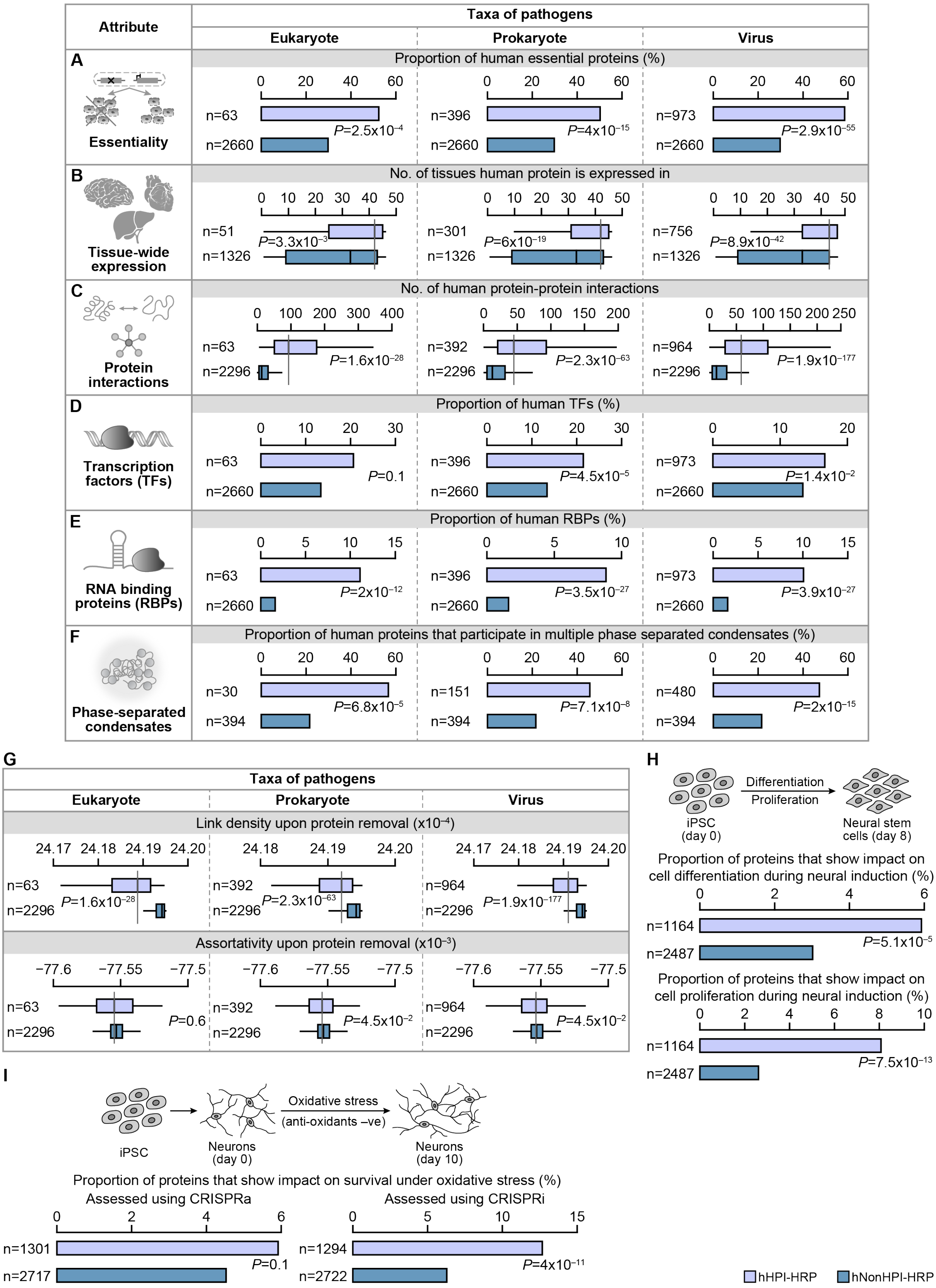


**Figure S2. Human HRPs engaged in HPI are physiological more important than human HRPs not engaged in HPI. (A)** Bar plots showing the proportion of essential human proteins with homorepeats. P value was estimated using Fisher's Exact test. Box plots representing the

**(B)** total number of tissues in which the different classes of human proteins are expressed and **(C)** number of protein-protein interactions in humans. P value was computed using Wilcoxon rank sum test. Bar plots showing the proportion of human **(D)** transcription factors (TFs) and **(E)** RNA binding proteins (RBPs) that are engaged by different pathogens in HPI. P value was computed using Fisher's Exact test. **(F)** Bar plots showing the proportion of human proteins that participate in more than one phase separated condensates. **(G)** Box plots representing the link density[7] (top panel) and assortativity[8] (bottom panel) of human proteins with homorepeats. P value was computed using Wilcoxon rank sum test. **(H)** Bar plots showing the proportion of proteins that show impact on cell differentiation (top panel) and cell proliferation (bottom panel) during neural induction of human iPSCs. **(I)** Bar plots showing the proportion of proteins that show impact on survival of neurons in oxidative stress assessed using CRISPRa (left panel) and CRISPRi (right panel). P value was estimated using Fisher's exact test. n denotes the total number of proteins in each class.

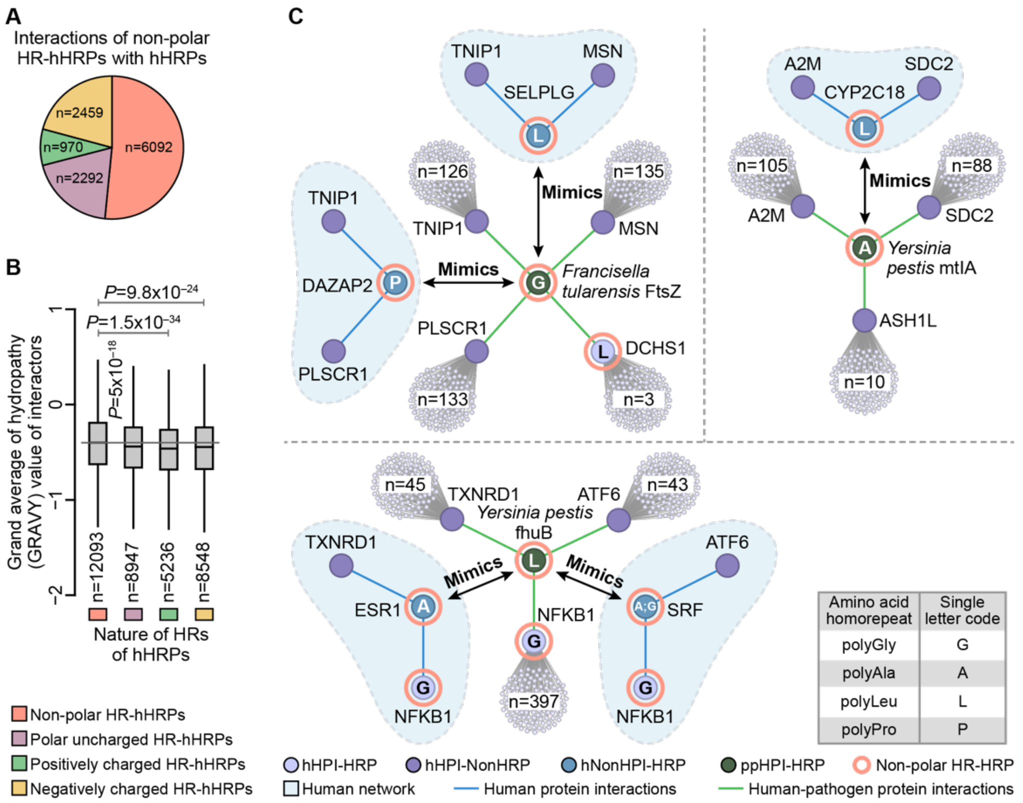


**Figure S3. hHRPs and pHRPs with non-polar HRs might interact with human proteins via hydrophobic interactions. (A)** Pie chart showing the total fraction of protein interactions of hHRPs with non-polar HRs with hHRPs with different physicochemical nature of HRs. n indicates the total number of protein interactions of hHRPs with non-polar HRs with each class of hHRPs. **(B)** Boxplot showing the hydropathy (GRAVY value; https://www.gravy-calculator.de/index.php) of proteins which are interactors of hHRPs with different physicochemical nature of HRs. P value was estimated using Wilcoxon rank sum test. n denotes the number of unique protein interactors of different classes of hHRPs. hHRPs with multiple HRs were considered only when all HRs were of the same physico-chemical nature. **(C)** Illustration showing instances of host protein proxy of ppHPI-HRPs with non-polar HRs. The node colour denotes the class of protein. The blue background colour denotes the human-physiological protein-protein interactions. Proteins with non-polar HRs are highlighted by a coral outline, with the type of HR denoted inside the node. n represents the number of direct protein interactors (first degree interactions) of hHPI proteins that interact with ppHPI-HRPs. ppHPI-HRP mimics hNonHPI-HRPs with non-polar HRs, thereby interacting with hHPI proteins, and subsequently influencing the first-degree interactors of hHPI proteins.

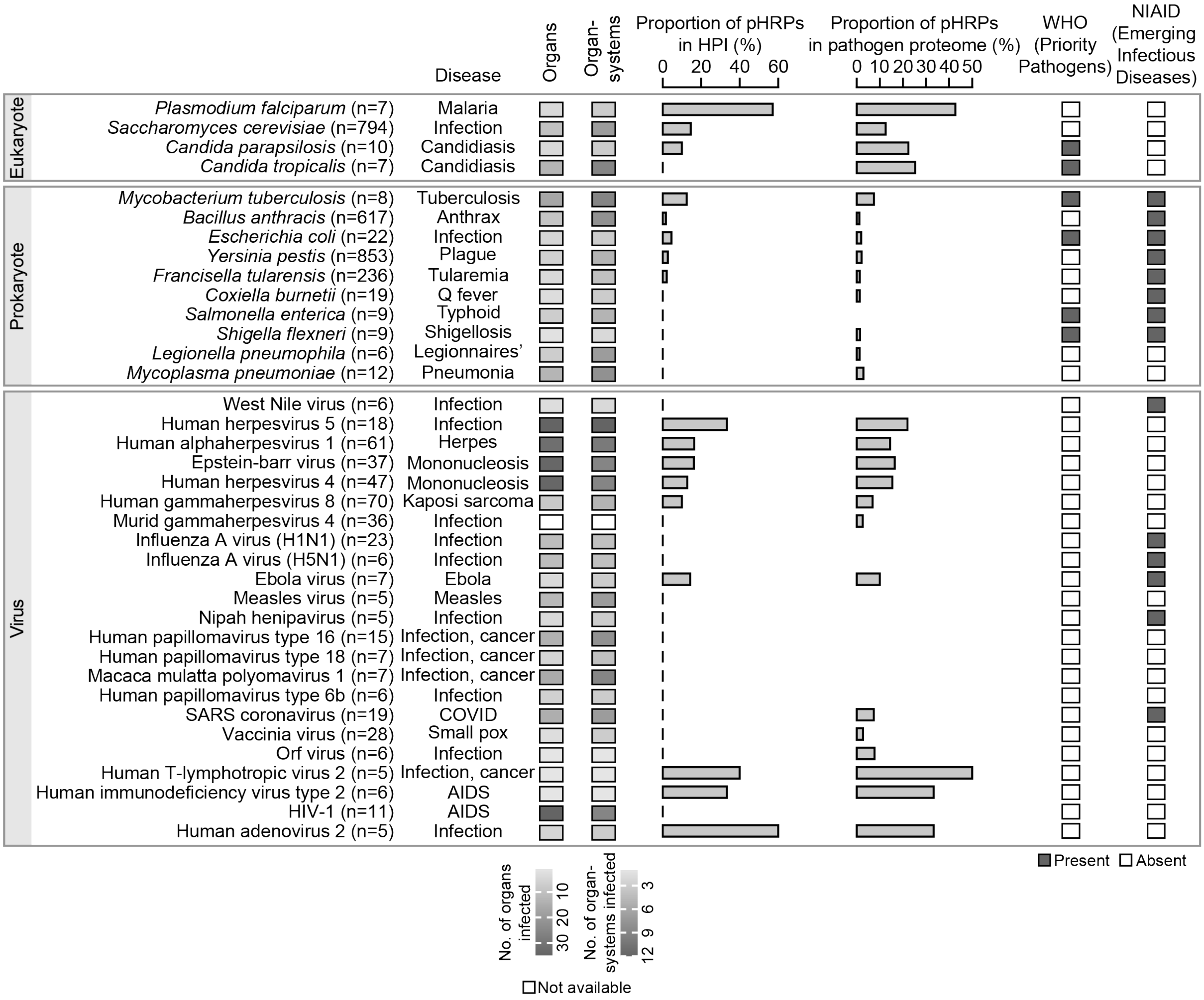


**Figure S4. Pathogens causing life-threatening diseases have high HRPs in HPI and their proteomes.** Integrated figure with (i) the heatmap showing the number of organs, organ systems affected by the pathogens, (ii) bar plots showing the number of pHRPs in HPI, pathogen proteome and classification of the pathogen as (iii) priority pathogens by World Health Organization (WHO) (iv) biodefense pathogens by National Institute of Allergy and Infectious Diseases (NIAID) (left to right). n denotes the number of pathogen proteins participating in HPI.

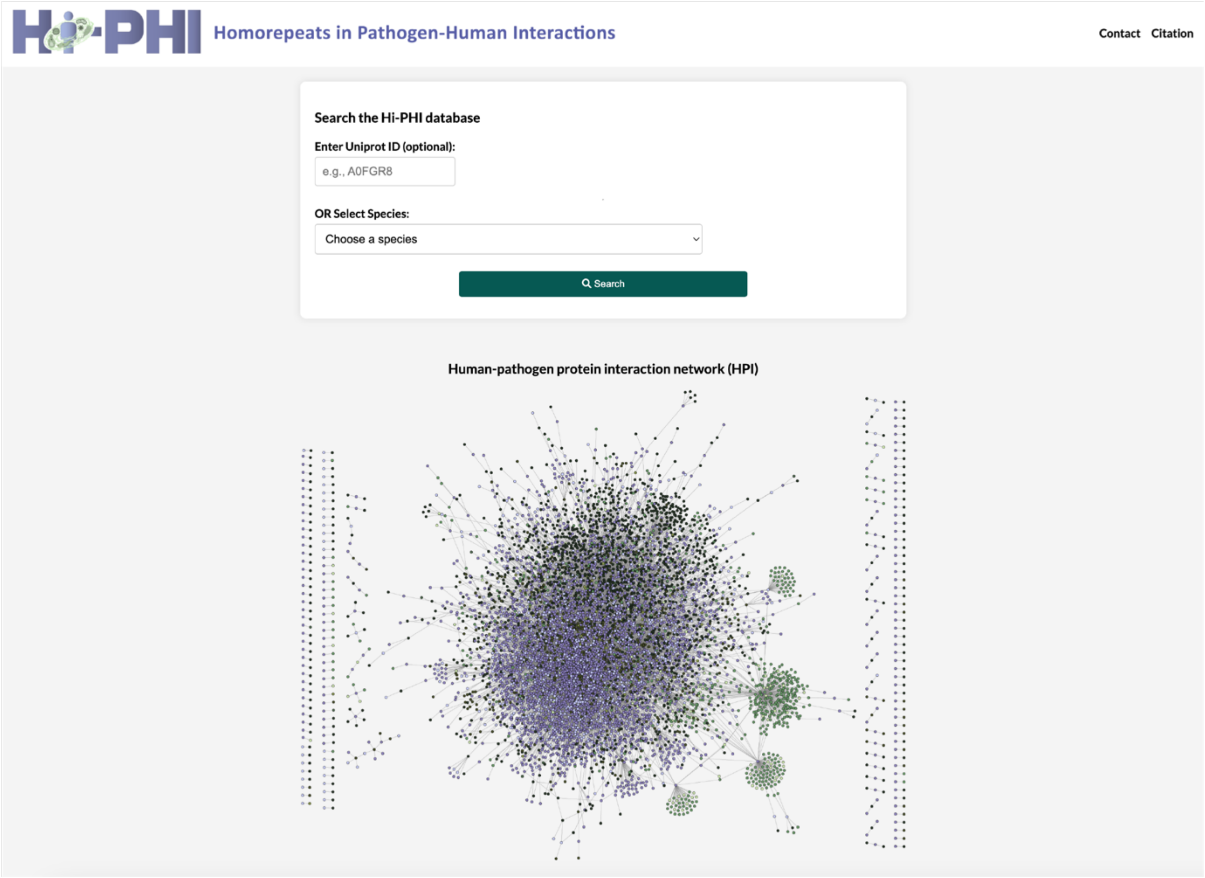


**Figure S5. Homorepeats in Pathogen-Human Interactions (Hi-PHI) resource.** Screen shot of the homepage of the database.
